# RocS drives chromosome segregation and nucleoid occlusion in *Streptococcus pneumoniae*

**DOI:** 10.1101/359943

**Authors:** Chryslène Mercy, Jean-Pierre Lavergne, Jelle Slager, Adrien Ducret, Pierre Simon Garcia, Marie-Francoise Noirot-Gros, Nelly Dubarry, Julien Nourikyan, Jan-Willem Veening, Christophe Grangeasse

## Abstract

Segregation of replicated chromosomes in bacteria is poorly understood outside some prominent model strains and even less is known about how it is coordinated with other cellular processes. Here we report that RocS is crucial for chromosome segregation in the opportunistic human pathogen *Streptococcus pneumoniae*. RocS is membrane-bound and interacts both with DNA and the chromosome partitioning protein ParB to properly segregate the origin of replication region to new daughter cells. In addition, we show that RocS interacts with the tyrosine-autokinase CpsD required for polysaccharide capsule biogenesis, which is crucial for *S. pneumoniae*’s ability to prevent host immune detection. Altering the RocS-CpsD interaction drastically hinders chromosome partitioning and cell division. Altogether, this work reveals that RocS is the cornerstone of an atypical nucleoid occlusion system ensuring proper cell division in coordination with the biogenesis of a protective capsular layer.

## Introduction

In dividing cells, accurate and faithful duplication and distribution of the genetic heritage are crucial steps toward the generation of viable and identical progeny. Unlike eukaryotes, in which chromosome segregation is performed by the well-known mitotic spindle (1), far less is known about the structures and mechanisms employed in bacteria. Intensive investigations have partly elucidated these processes in model bacteria like *Escherichia coli*, *Bacillus subtilis* and *Caulobacter crescentus* (for reviews see (2-5). However, these studies also highlight that there are large variations in the mechanisms involved, and it remains poorly understood in many other bacterial species.

This is the case for *Streptococci* and more specifically the opportunistic human pathogen *Streptococcus pneumoniae* (the pneumococcus), a prominent model to study the bacterial cell cycle (6). The pneumococcus is an ovoid-shaped bacterium that lacks some of the well-established systems controlling cell division, like the Min and the nucleoid occlusion systems (7). It possesses only the condensin SMC and an incomplete chromosome partitioning ParABS system, in which ParA is absent. Previous studies have evidenced that both ParB and SMC are involved, but not essential, in pneumococcal chromosome segregation (8). Notably, both individual or double deletion of *parB* and *smc* only lead to weak chromosome segregation defects, suggesting that other factors remain to be discovered. In line with this hypothesis, transcription was shown to contribute to pneumococcal chromosome segregation (9). The tyrosine-autokinase CpsD was also found to interfere with chromosome segregation (10). CpsD is primarily described as a key regulator of the export and synthesis of the polysaccharide capsule, the main virulence factor of the pneumococcus, that is exclusively produced at the pneumococcal division septum (10-13). However, defective autophosphorylation of CpsD also generated elongated cells with an aberrant nucleoid morphology (10).

## Results and Discussion

To further analyze the relationship between capsule production and chromosome biology, we screened a yeast two-hybrid genomic library of a pneumococcal laboratory strain (14) using CpsD as bait. A strong and reproducible interaction was identified with Spr0895, a protein with unknown function (Fig. S1A). The CpsD-Spr0895 interaction was confirmed biochemically using microscale thermophoresis (Fig. S1B-C). The *spr0895* gene is conserved among *Streptococcaceae* (Fig. S2) and is hereinafter referred to as *rocS* (<u>R</u>egulator <u>o</u>f <u>C</u>hromosome <u>S</u>egregation) based on the observations we report below.

We first constructed a *rocS* deletion in the encapsulated virulent D39 strain and analyzed capsule production by immunofluorescence microscopy using anti-serotype 2 capsule antibodies (10). As observed for wild type cells, capsule was detected over the entire surface of Δ*rocS* cells (Fig. 1A). In addition, immunodetection of the total fraction of capsule by western-blot revealed that capsule production and polymerization were not affected (Fig. S3). However, although the cell shape of Δ*rocS* cells was not significantly altered, they displayed a growth defect with an increased generation time compared to wild type cells (Fig.S4). Surprisingly, when we looked at the DNA content of Δ*rocS* cells using DAPI staining, we found that 13.9% of cells were anucleate (Fig. 1A-B). Unencapsulated and non-virulent laboratory R800 cells deficient for *rocS* showed similar growth defects and a comparable fraction of anucleate cells (15.7%), indicating that these aberrant phenotypes were not dependent on capsule production (Fig. 1B and S5). Complementation of the ΔrocS in both R800 and D39 genetic backgrounds with an ectopic copy of *rocS* (Δ*rocS*-P_comX_-*rocS*) restored the wild type phenotype with 1% and 1.5% of anucleate cells, respectively (Fig. 1B). By comparison, the deletion of well-established factors required for chromosome replication and/or segregation like ParB or SMC in *S. pneumoniae*, results in less than 4% and 2% of anucleate cells, respectively (10). Furthermore, we were unable to delete both *rocS* and either smc or *parB*, indicating that the deletion of *rocS* is synthetically lethal with *parB* or *smC*. Altogether, our results show that RocS has an important role in pneumococcal chromosome biology.

**Fig. 1:**
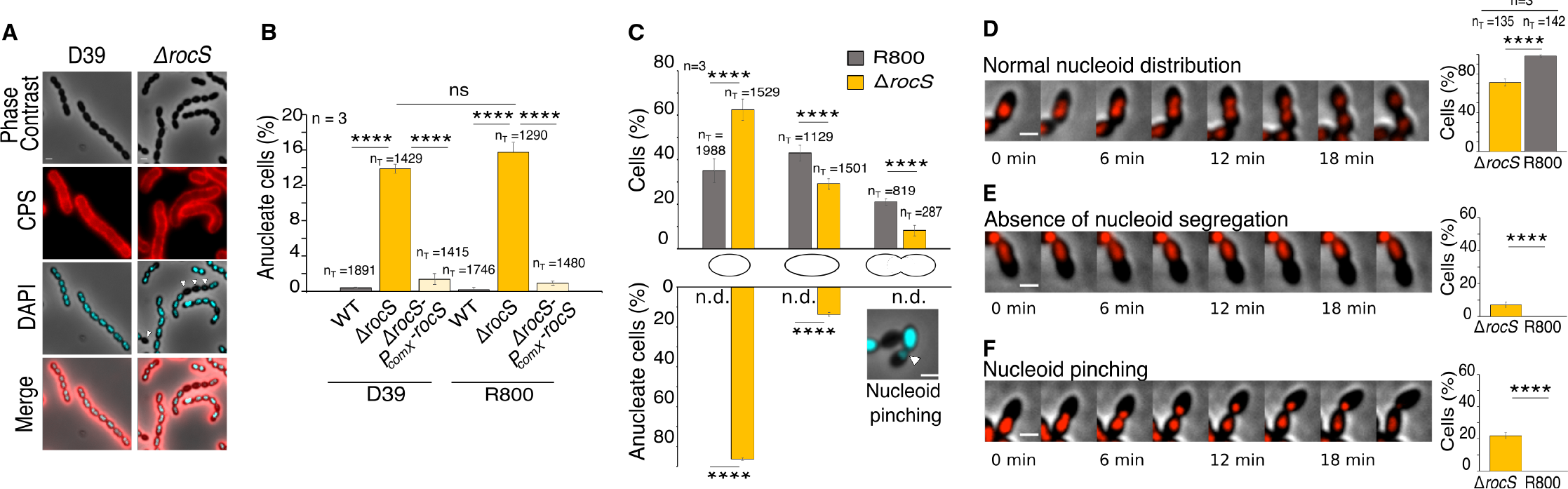
Impact of *rocS* deletion on capsule production and nucleoid distribution. (A) Detection of capsular polysaccharides (CPS) and DNA in D39 and Δ*rocS* cells. Phase contrast (grey), CPS (red), DAPI (blue) and overlays are shown. Arrow heads indicate anucleate cells. (B) Percentage of anucleate cells in D39 and R800 (grey) strains, corresponding Δ*rocS* mutants (orange) and complemented strains (yellow). (C) Quantification of anucleate cells in the course of the cell cycle. R800 (grey) and Δ*rocS* (orange) cells were classified into three groups (nascent, elongated, constricting) according to the progression of the cell cycle. The top histogram shows the percentage of each group for each strain. The percentage of anucleate cells in each group is shown in the bottom histogram. Arrowheads indicate chromosome pinching in constricting cells. n.d.= none detected. In (B) and (C), standard deviations are indicated with error bars. (D-F) Still images from fluorescence time-lapse microscopy (Movies S1, S2 and S3) of WT (D) and Δ*rocS* cells (E and F) producing HlpA-mKate2. (D) Normal nucleoid segregation, (E) absence of nucleoid segregation and (F) nucleoid pinching. Histograms show the percentage of the nucleoid organization (absence or pinching) in WT and Δ*rocS* cells. In B-F, nT indicates the number of cells analyzed from 3 independent experiments and standard errors are indicated with error bars. (Two population proportions test: **** P < 0.0001. ns P > 0.5). Scale bar, 1 μm.

To understand the function of RocS, we analyzed Δ*rocS* R800 cells at three different stages of the cell cycle (nascent, elongated and constricted cells, (Fig. 1C). By comparison with the relative proportion observed for wild type cells, we observed a significant increase of cells at the early stage of the cell cycle, relative to cells at the later stages: 62.5% of *ΔrocS* cells displayed the typical morphology of rounded nascent cells while only 35% of wild-type cells harbored this morphology (Fig. 1C). Strikingly, a large majority of anucleate cells (86.3%) were indeed observed at the early stage of the cell cycle (nascent cells). In addition, some cells at the later stages (constricted cells) of the cell cycle harbored asymmetric distribution of the DNA content, which suggests chromosome-pinching events (Fig. 1C).

To get more insight into chromosome dynamics in the absence of RocS, we performed time-lapse microscopy to image the nucleoid by localizing the pneumococcal histone-like protein, using a HlpA-mKate2 fusion (9). As expected, the chromosome duplicates at the early stage of the cell cycle and eventually splits into two parts that segregate to each daughter cell (Fig. 1D and Movie S1). In contrast, newly replicated chromosomes in Δ*rocS* cells were either not segregated (7%) (Fig. 1E and Movie S2), or partially segregated and eventually truncated by the newly forming septum (21.8%), a process also known as the guillotine effect (15) (Fig. 1F and Movie S3). In the latter case, the signal of the truncated chromosome became diffuse and was ultimately degraded. In both cases, these aberrant chromosome-partitioning events led to the formation of anucleate nascent cells. To test if chromosome replication was affected in the R800 Δ*rocS* mutant, we used qPCR to determine the ratio between the origin of replication (*oriC*) and the terminus region (*ter*) of the chromosome in exponentially growing cells (16) (Fig. S6). As expected, we observed that dividing wild type cells displayed a characteristic mean ratio of 1.68 ±0.28 whereas this ratio was close to 1 for a thermo-sensitive *dnaA* (the replication initiator protein) mutant shifted to non-permissive temperature. The origin-to-terminus ratios of *ΔrocS* (1.67 ± 0.24) and complemented *ΔrocS-P_comX_-rocS* (1.56 ± 0.24) cells were similar to that of wild type cells, indicating that RocS is not involved in chromosome replication. Together, our results show that chromosome segregation rather than chromosome replication is severely affected in the absence of RocS.

To characterize the contribution of RocS to chromosome segregation, we next examined the subcellular localization of the origin of replication (*oriC)* during the cell cycle of wild-type and Δ*rocS* R800 cells (Fig. 2A-D). To visualize *oriC* at the single cell level, we engineered a system based on the ectopic production of a fluorescent fusion of RepC, the ParB homolog of *Enterococcus faecalis,* and insertion of *parS_Ef_* sites from *E. faecalis* near the pneumococcal *oriC* (17) (Fig. 2A). Neither expression of *repC-gfp* nor insertion of *parS_Ef_* sites influenced the pneumococcal cell cycle as evidenced by wild-type growth kinetics and cell morphology (Fig. S7). When produced, the RepC-GFP fusion, which binds specifically to the *parS_Ef_* site, formed diffraction-limited foci in the vicinity of *oriC,* (Fig. 2B and Fig. S7). As previously characterized (18), *oriC* localized as a single focus located around mid-cell of nascent cells (Fig. 2B). The duplication of the focus was followed by rapid segregation of the two foci toward the center of each daughter cell where they remain as the cell elongate. Interestingly, new cycles of chromosome segregation started early in the cell cycle, even before the completion of division, as attested by the 4.5% of nascent cells containing 2 foci and the 5% of cells at the later stage of the cell cycle containing 3 or 4 foci (Fig. 2B and 2D). By comparison, the subcellular localization of *oriC* throughout the cell cycle was strongly affected in the absence of RocS. After duplication, most of the two foci remained near mid-cell and did not segregate toward the opposite poles as the cells elongate (Fig. 2C). On average, the spacing rate (relative distance between 2 foci of *oriC* in relation to the cell length) was significantly lower in *ΔrocS* cells (0.32±0.003) than in WT cells (0.47±0.003) (Fig. 2E). Furthermore, the proportion of cells with single foci was significantly higher in *ΔrocS* cells (47.6%) than in wild-type cells (23%). Since chromosome replication was not affected in *ΔrocS* cells (Fig. S6), this observation suggests that after replication, some *oriC* copies may be too close to be detected as separated foci in *ΔrocS* cells. Finally, we did not detect constricting cells containing 3 or 4 foci in *ΔrocS* cells (Fig. 2C and 2D). Altogether, these data show that the two newly replicated chromosome origins segregate less efficiently in the absence of RocS, reflecting its crucial role in chromosome segregation.

**Fig. 2:**
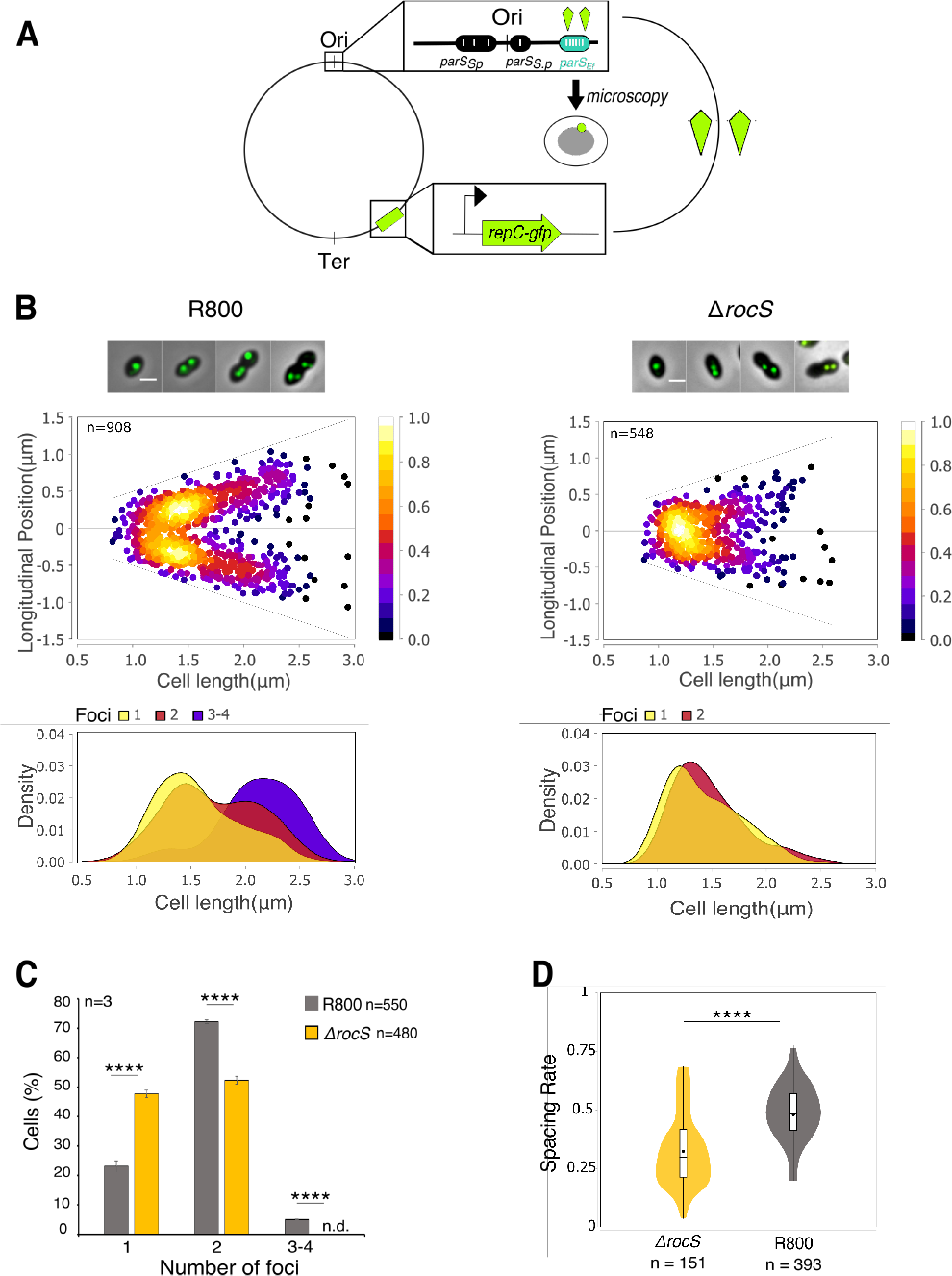
*oriC* segregation patterns in wild-type and *ΔrocS* cells. (A) Schematic representation of the Par system used to image the origin of replication (*oriC*). ***parS*** sequences from *E. faecalis* (*parS_Ef_*, blue oval) were inserted into the chromosome near the pneumococcal *oriC* while the *parB* homolog *repC* fused to *gfp* (RepC-GFP, green kite) is expressed ectopically under the control of the *P_comX_* promoter. Upon loading of GFP-RepC onto parS*_Ef_* sites, the localization of *oriC* is followed by fluorescence microscopy (green dot). *parS_Sp_* indicates native pneumococcal *parS* sites. (B) (upper panels) Localization heat maps of *oriC* (GFP-RepC) positions along the cell length in wild-type and Δ*rocS* R800 cells. Representative overlays between phase contrast and GFP fluorescence signal of cells with either 1, 2 or 3/4 foci are shown on the top. Scale bar, 1 μm. (lower panels) Kernel density plots of the cell length in relation to the number of foci in wild-type and *ΔrocS* R800 cells. (C). Quantification of cells as a function of the number of *oriC* foci in WT (grey) and *ΔrocS* (orange) cells. Standard errors are indicated with error bars. (D). Measurements of the spacing rate (relative distance between 2 foci of *oriC* in relation to the cell length). nT indicates the number of cells analyzed. Experiments were performed in triplicates. (Two population proportions test: **** P < 0.0001).

We next followed the subcellular localization of RocS during the cell cycle using a reporter strain expressing a fluorescent fusion of RocS. The gene *gfp-rocS* replaces the *rocS* gene at the native chromosomal locus and encodes a largely functional fusion protein, as attested by wild-type growth kinetics, cell morphology, intracellular level and low level of anucleate R800 cells (3%) (Fig. S8 and S9).

The GFP-RocS fusion protein formed faint diffraction-limited foci that were mobile, able to both fuse into stationary and brighter foci and to re-segregate (Movie S4). Interestingly, while mobile foci showed no specific localization during the cell cycle, the brighter foci showed a dynamic reminiscent of that of *oriC. I*ndeed, brighter foci mostly localized around mid-cell of nascent cells and positioned toward the center of the daughter cell as the cell elongates (Fig. 3A). Bioinformatic analysis of the RocS sequence predicted the presence of a C-terminal membrane-binding amphipathic helix (AH) homologous to that of MinD of *Escherichia coli* (19) and an N-terminal helix-turn-helix domain (HTH, InterPro IPR000047) characteristic of DNA-binding proteins (20) (Fig. S10). These two domains are required for the function of RocS in chromosome segregation as both *ΔHTH-rocS* and *rocS-ΔAH* R800 cells displayed growth and viability defects as well as an anucleate phenotype similar to *ΔrocS* R800 cells (Fig. 3D and Fig. S11). In addition, deletion of either the AH or the HTH domains drastically altered the localization pattern of RocS (Fig. 3B-C). The deletion of the N-terminal HTH domain resulted in the discontinuous redistribution of GFP-ΔHTH-RocS at the cell periphery. On the other hand, GFP-RocS-ΔAH displayed a diffused localization in the pneumococcal cell, which co-localized with the nucleoid (median R = 0.85, interquartile range = 0.83-0.92) (Fig. 3C). Using gel shift assays, we showed that RocS-ΔAH binds directly to DNA (Fig. S12). This DNA binding was independent of the length, the GC content or the sequence of the tested DNA. Altogether, these data show that the C-terminal AH is required for the interaction of RocS with the membrane, while the N-terminal HTH domain mediates RocS DNA binding. Collectively, these data show that the interactions of RocS with both the chromosome and the membrane are essential for its function in chromosome segregation.

**Fig. 3.**
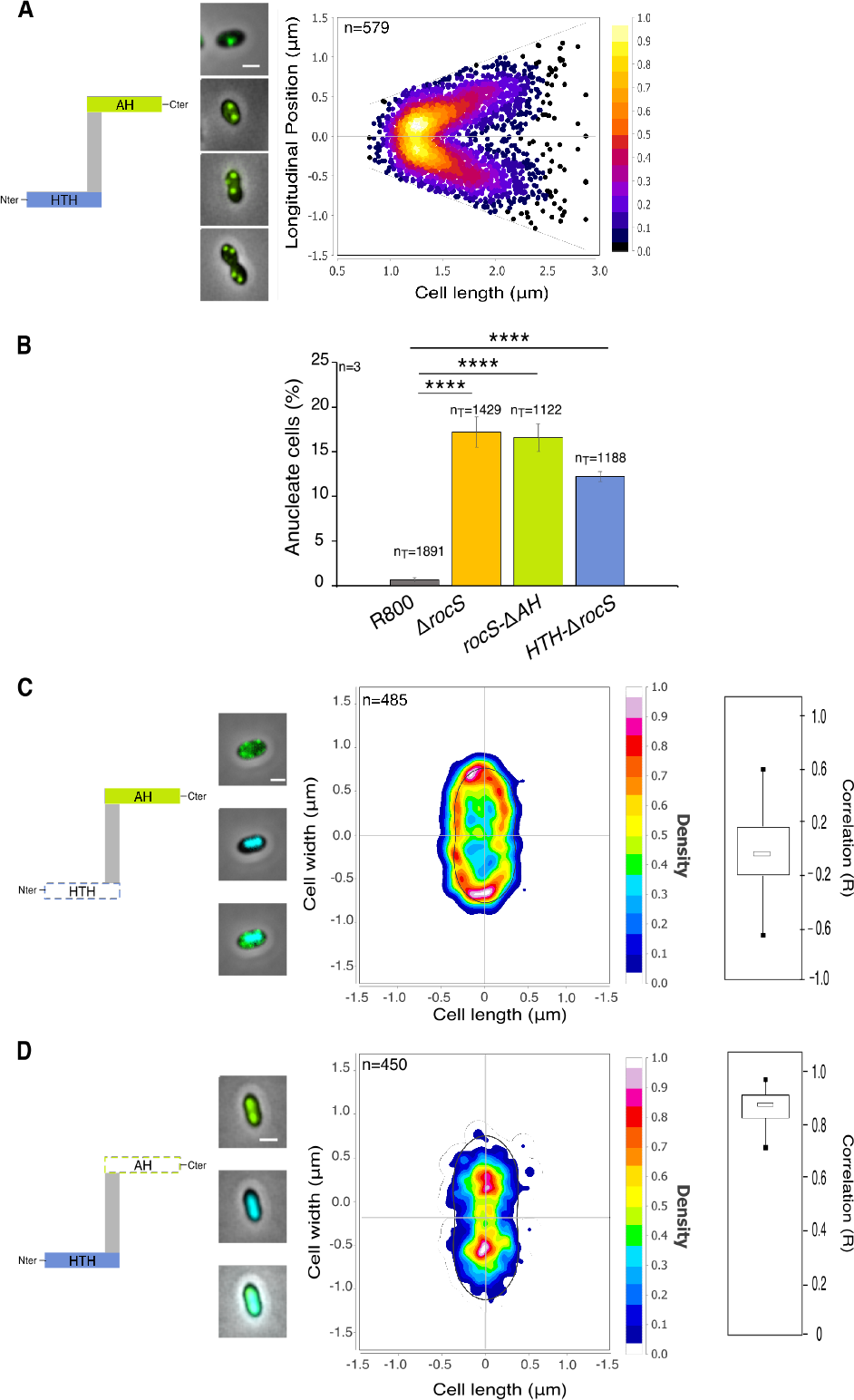
Localization of GFP-RocS and derivatives and impact on nucleoid localization. Schematic representations of RocS and derivatives are shown on the left of panels A, C and D. (A) Heat map representing the longitudinal localization of GFP-RocS as a function of the cell length in R800 cells. Representative overlays of cells with either 1, 2 or 3/4 foci are shown on the left. (B) Histogram showing the percentage of anucleate cells for *rocS-ΔAH* and *ΔHTH-rocS* R800 strains. Standard errors are indicated with error bars. n_T_ indicates the total number of cells analyzed from three independent experiments. (Two population proportions test: **** P<0.0001). (C-D) Heat map representing the 2-dimensional localization patterns of GFP-ΔHTH-RocS (C) and GFP-RocS-*Δ*AH (D) in R800 cells. Representative overlays of phase contrasts and, GFP or DAPI fluorescence signals, or both signals, are shown on the left of the heat maps. Scale bar, 1 μm. The distribution of the Pearson correlation coefficient (R), measured between the DAPI and GFP signals for each strain are shown as box and whisker plots on the right.

We finally questioned the biological role of the interaction between RocS and the tyrosine-autokinase CpsD (Fig. S1). Previous findings showed that CpsD possesses a structural fold comparable to that of ParA proteins that usually assist ParB in chromosome segregation (10, (21, 22). Since ParA is absent in the pneumococcus (7) and CpsD interacts directly with ParB, it was proposed that CpsD could act as a ParA-like protein (10). Interestingly, this interaction is modulated by the autophosphorylation of CpsD: mimicking permanent phosphorylation of CpsD(CpsD-3YE) promotes capsule biogenesis and normal chromosome segregation by enabling ParB mobility (10) (Fig. 4A). By contrast, defective autophosphorylation of CpsD (CpsD-3YF) not only impairs capsule production, but also reduces ParB mobility, inducing aberrant chromosome segregation and leading to cell elongation (10) (Fig. 4B). By consequence, even in the absence of a conserved nucleoid occlusion system in the pneumococcus (7), cell division seems inhibited to protect the nucleoid against truncation by the newly forming septum when CpsD is not phosphorylated. Interestingly, we also demonstrated that RocS interacts with ParB both *in vivo* and *in vitro* (Fig. S13). Therefore, we wondered if this cell division block was due to RocS. To test this hypothesis, we deleted *rocS* in D39 strains mimicking either permanent or defective phosphorylation of CpsD (respectively Δ*rocS-cpsD-3YE* and Δ*rocS-cpsD-3YF*) and looked at the cell morphology, capsule production and DNA content. While deletion of *rocS* in the permanent phosphorylation *cpsD-3YE* mutant did not impact the cell morphology, the deletion of *rocS* suppressed the elongated phenotype of the defective phosphorylation *cpsD-3YF* mutant (Fig. 4A). In both cases, the deletion of *rocS* is accompanied by approximately 13% of anucleate cells (12.8% and 13.7% respectively). The deletion of *rocS* in the phospho-ablative mutant therefore abrogates the cell division block. Non-phosphorylated CpsD together with RocS could therefore be viewed as a *bona fide* nucleoid occlusion system.

**Fig. 4.**
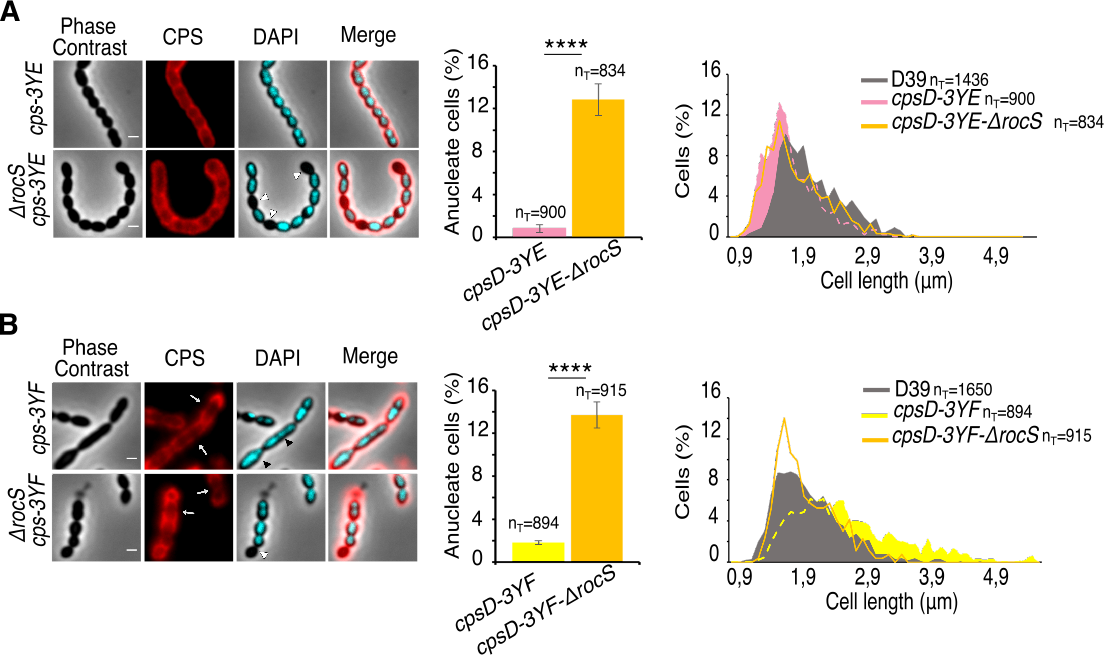
Deletion of *rocS* in phospho-ablative and phospho-mimetic CpsD mutants. Detection of CPS and DNA in (a) *cpsD-3YE* and *cpsD-3YE-ΔrocS* and (b) *cpsD-3YF* and *cpsD-3YF-ΔrocS*. Phase contrast (grey), CPS (red), DAPI (blue) and overlays are shown on the left. White arrows show CPS production defects, white arrowheads show anucleate cells and black arrowheads show nucleoid segregation defects. Scale bar, 1 µm. The corresponding percentage of anucleate cells are shown in the middle. (Two population proportions test: **** P < 0.0001). The corresponding distribution of the cell length are shown on the right. nT indicates the number of cells analyzed from 3 independent experiments and standard errors are indicated with error bars.

Typical nucleoid occlusion systems prevent the assembly of the FtsZ ring over the nucleoid. However, the FtsZ-ring is properly positioned in elongated cells (10) indicating that cell constriction, but not FtsZ-ring assembly, is blocked in the *cpsD* phospho-ablative mutant. As the deletion of *rocS* alleviates this phenotype, one can further conclude that RocS is the cornerstone of a new type of nucleoid occlusion system that prevents the constriction of the FtsZ-ring rather than impeding its assembly (Fig. 5). Which exact step of cell septation is blocked by the here-identified nucleoid occlusion system remains unclear, but it is tempting to speculate that RocS could work in partnership with the major cell division regulator of the pneumococcus, the eukaryotic-like protein-kinase StkP, whose inactivation also results in cell elongation. (23, 24). In any case, this work illustrates that pathways controlling chromosome segregation and cell division are far more diverse than expected and suggests that the RocS system is likely valid for all *Streptococcaceae* (Fig. S2). The "raison d'être" of such a regulatory process coordinating capsule synthesis with cell cycle progression is likely to make sure that cells are covered by capsule at every step of the cell cycle in order to prevent detection by the human immune system.

**Figure 5:**
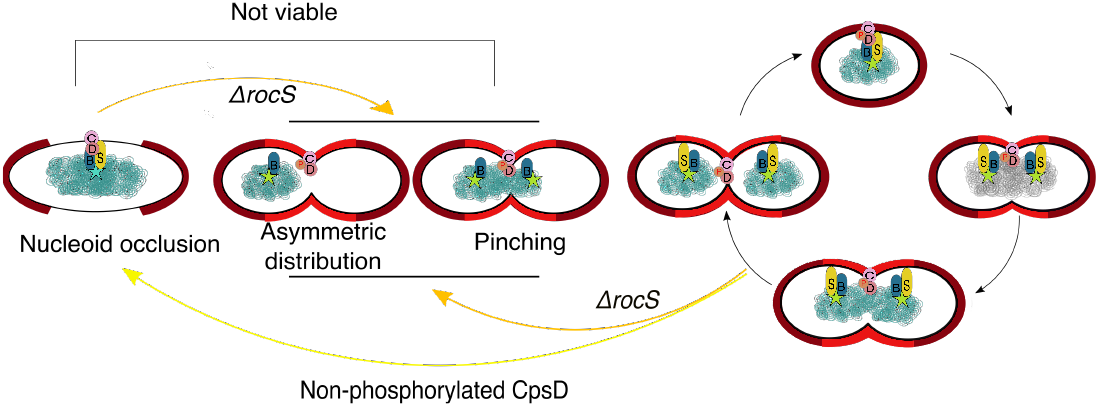
Model for the nucleoid occlusion system coordinating capsule synthesis, chromosome segregation and pneumococcal cell division. When CpsD is autophosphorylated, capsule is properly produced and RocS and ParB actively contribute to chromosome segregation (viable encapsulated progeny). By contrast, non-phosphorylated CpsD hinders both capsule synthesis and chromosome segregation inducing a division block. The deletion of *rocS* alleviates the division block and results in uncontrolled cell constriction with improper chromosome segregation (pinching and asymmetric distribution) leading to non-viable progeny. ParB, RocS, CpsD and its transmembrane activator CpsC are indicated by blue, yellow, brown and pink circles, respectively. Red "P" and the turquoise star indicate CpsD autophosphorylation and the *oriC* region, respectively. Capsule is shown in light (new capsule produced during cell division) and dark (inherit from the mother cell) red.

## Materials and Methods

### Strains and growth conditions

Strains used in this study are listed in the Table S1. *Streptococcus pneumoniae* R800 and D39 and derivatives were cultivated at 37°C in C+Y medium or Todd-Hewitt Yeast (THY) broth.

Cell growth curves were monitored in JASCO V-630-BIO-spectrophotometer and the optical density was read automatically every 10 min. *Escherichia coli* XL1-B strain (25) was used for cloning and *E. coli* BL21 (26) for overproduction of CpsC/D, RocS, RocS-ΔAH and ParB. *E. coli* strains were grown in Luria Bertani broth (LB) supplemented with appropriate antibiotic. Growth was monitored by optical density (OD) readings at 550 nm or 600 nm for *S. pneumoniae* or *E. coli* strains, respectively.

### Construction of plasmids and strains

Gene modifications (*gfp* and *flag* fusions, knock-out and domain deletion) in *S. pneumoniae* were achieved by homologous recombination using the two-step procedure based on a bicistronic *kan-rpsL* cassette called Janus (27) and constructed at their native chromosomal locus. They are thus expressed under the control of the native promoter and represent the only source of protein.

Δ*rocS* D39 and Δ*rocS* R800 strains were complemented ectopically for *rocS* expression using the strategy described by (28) using the competence inducible system of *Streptococcus thermophilus*. The ComS-inducible *comR* DNA fragment was introduced between the *treR* and *amiF* loci of both strains. Then, the *rocS* copy under the control of the *comX* promoter was inserted between the *cpsN* and *cpsO* genes in R800 or at the *bgaA* locus in D39 strains.

For constructing the system for tagging *ori*, we used the *parS* sites and the ParB homologue RepC fused to the GFP from *Enteroccocus faecalis* (17). The *parS* sites were inserted between *thmA* and IS1167 loci near the pneumococcal origin of replication. Then, the *repC-gfp* under the control of the promoter of the *comX* gene of *Streptococcus thermophilus* were used by PCR and inserted between the *cpsN* and *cpsO* genes in the R800 strain. *repC-gfp* expression was induced with 5 µM ComS.

To construct the thermo-sensitive *dnaA* R800 mutated strain, we PCR amplified the *dnaA(T1193C)* mutated gene of the D39 thermo-sensitive mutant described in (9). The DNA fragment was then transformed in the R800 strain and cells were plated at 30°C. After overnight growth, colonies were resuspended in THY and cultured again on plates at either 30°C or 40°C. The mutation in *dnaA* was checked by DNA sequencing in clones growing at 30°C but not at 40°C.

For the construction of the plasmid overexpressing RocS-ΔAH-6His, we PCR amplified a DNA fragment coding for RocS from Met1 to Gln150 using chromosomal DNA from *S. pneumoniae* R800 as template. The obtained DNA fragment was cloned between the *NdeI* and *PstI* cloning sites of pT7-7 (29). The other plasmids used in this study are described in Table S1.

The oligonucleotides used for all construction are listed in Table S2. Plasmids and pneumococcal strains were verified by DNA sequencing to verify error-free PCR amplification.

### Protein purification

Purification of the chimera 6His-CpsC/D and ParB-6His was performed as described previously (10). To purify RocS-ΔAH-6His, *E. coli BL21* were used and cultured at 37 °C in LB medium. At OD_600_ = 0.6, 1mM IPTG was added and ells culture were incubation was continued for 3 h. at 37 °C. Cells were then harvested by centrifugation and resuspended in buffer A (Tris-Hcl 25 mM, pH 7.5; NaCl 1 M, imidazole 10 mM; glycerol 10%) containing 10 mg mL^-1^ of lysozyme, 1 µg ml^−1^ of protease inhibitor (Roche Diagnostics). After sonication and centrifugation, the supernatant was loaded on to a Ni-NTA agarose resin (Qiagen) and extensively washed with buffer A containing 20 mM imidazole. RocS-6His was eluted with buffer B (Tris-Hcl 25 mM, pH 7.5; NaCl 300 mM, imidazole 300 mM; glycerol 10%). Pure fractions were pooled and dialyzed against buffer C (HEPES 50 mM, pH 7.5 or Tris pH 7,5 25 mM; NaCl 150 mM, glycerol 10%). Protein concentrations was determined using a Coomassie assay protein dosage reagent (Uptima).

### Co-immunoprecipitation and immunoblot analysis

For co-immunoprecipitation, cultures of *S. pneumoniae* cells were grown at 37°C in C+Y medium until OD_550nm_ = 0.3. Cells pellets were incubated at 30°C for 30 min in buffer A (Tris-HCl 0.1 M, pH 7.5; MgCl_2_ 2 mM, Sucrose 1 M, 6 mg mL^-1^ of DNase I and RNase A, 1 µg ml^−1^ of protease inhibitor). After centrifugation at 4°C, the pellet was resuspended in buffer B (Tris-Hcl 0.1 M, pH 7.5; EDTA 1 mM, 0.1% Triton, 6 mg mL^-1^ of DNase I and RNase A, 1 µg ml^−1^ of protease inhibitor) and incubated 15 min at room temperature before being harvested by centrifugation. The supernatant was then incubated with Dynabeads (Invitrogen) coupled with 20 µg anti-Flag antibodies and incubated for 2 hour at 4°C. After extensive wash with buffer C (Tris-Hcl 10 mM, pH 7.5, EDTA 0.5 mM, 0.1% Triton, NaCl 150 mM, 1 µg ml^−1^ of protease inhibitor), Protein-bounded bead were eluted with SDS-PAGE loading buffer at 95°C for 10 min and analyzed by SDS-PAGE and immunoblotting using a rabbit anti-GFP antibody at 1/10,000 (AMS Biotechnology) or the anti-FLAG antibody at 1/1,000 (Sigma). For immunoblot analysis, *S. pneumoniae* pellets were resuspended in TE-buffer (25 mM Tris-HCl pH 7.5, 1 mM EDTA) supplemented with protease and phosphatase inhibitor cocktail II (Sigma-Aldrich) and opened by sonication. 25 µg of crude extracts were analyzed by SDS-PAGE, electrotransferred onto a polyvinylidene difluoride membrane and incubated with either rabbit anti-RocS at 1/5,000 (produced by Eurogentec with purified RocS-ΔAH-6His), rabbit-anti-enolase polyclonal antibody at 1/50,0000 (23) or rabbit anti-serotype 2 CPS polyclonal antibody at 1/2,000 (Statens serum Institute). A goat anti-rabbit polyclonal antibody horseradish peroxidase (HRP) conjugated (Biorad) was used at 1:5000 to reveal immunoblots.

### Yeast-two hybrid

The yeast two hybrid genetic screens were carried out using a mating strategy as described previously (14) (30). Construction of the pGBDU-*cpsD* bait plasmid and expressing CpsD fused to the DNA-binding domain of Gal4 (BD) was described in (10). This plasmid was introduced in the PJ69-4(ɑ) haploid strain. This strain was then mated with PJ69-4 haploid(ɑ) strain harboring a library of pGAD plasmids expressing genomic fragments of *S. pneumonaie* R6 in fusion with the GAL4 activating domain (AD) (14). Potential binary interactions were selected by the ability of the yeast diploids to grow on synthetic media agar SC–LUH lacking Leucine (L) and Uracil (U) to select for maintenance of plasmids pGAD and pGBDU, respectively, as well as histidine (H), to selects for the interaction (31). Additionally, binary interactions were tested by a matrix-based approach by mating haploid cells expressing BD-CpsD, with haploid cells of complementary mating type expressing the AD-prey protein fusions RocS_50-163_, RocS, CpsC and CpsD. Diploids were first selected onto –LU media and further tested for interacting phenotypes (i.e. ability to grow on SC–LUH selective agar plates) to reveal binary interactions between bait and prey proteins.

### Preparation and analysis of CPS

CPS were prepared as previously described (10). Briefly, *S. pneumoniae* cultures were grown until OD_550nm_ = 0.3, washed once with PBS and resuspended in buffer A (Tris-HCL 50nM, pH 7.4; sucrose 20%; MgSO_4_ 50 nM). The solution was then added with 400 units of mutanolysin and 6 µg/µl of DNase and RNase and incubated overnight at room temperature. After centrifugation at 16,000 × g for 20 min at 4 °C, pellets were resuspended in the same volume of buffer A. 10 µL of the mixture were then mixed with 5 µl of buffer B (Tris-HCl 50 mM, pH 8.0; EDTA 50 mM; Tween20 0.5%; Triton X100 0.5%) and 20 µg of proteinase K, incubated 30 min at 37°C and analyzed by SDS-PAGE and immunoblotting.

### Microscopy techniques

Cells were grown until OD_550nm_ = 0.1. For immunofluorescence microscopy, cells were mixed with the rabbit-serotype 2 CPS polyclonal antibody (Statens Serum Institute) at 1/1,000, washed and then incubated with the anti-rabbit Dylight-549 antibody (Jackson ImmunoResearch) at 1/2,000. After a last wash with PBS, CPS were imaged.

For DAPI staining, 10 µl of *S. pneumoniae* cell culture were mixed with 1 µl of DAPI at 2 µg/µl (Molecular Probes) and incubated 5 min at room temperature. For mkate2 and GFP fluorescence imaging, cells were spotted on pads made of 1.5% agarose in C+Y medium at 37°C as described in (32). Slides were visualized with a Nikon TiE microscope fitted with an Orca-CMOS Flash4 V2 camera with a 100 × 1.45 objective. Images were collected using NIS-Elements (Nikon). Images were analyzed using the software ImageJ (http://rsb.info.nih.gov/ij/) and the plugin MicrobeJ (33).

Diffraction-limited foci of RepC-GFP or GFP-RocS were detected using the feature/spot detection option in MicrobeJ. This option combines spatial 2D filtering and 2D local maxima algorithm to localize single fluorescent maxima in each detected cell. Each maximum was then fit to a single peak or a multi peak 2D Gaussian curve, to determine their amplitude, their FWHM (Full width at half Maximum) and their coordinates at the subpixel resolution. Maxima were finally filtered based on the goodness of the fit and their amplitude. Their sub-cellular localizations were automatically computed for each associated particle.

### Microscale thermophoretic analysis

Microscale thermophoresis was used to test the interaction of RocS-AH with the chimeras CpsC/D and ParB (34). Binding experiments were carried out with a Monolith NT.115 Series instrument (Nano Temper Technologies GMBH). RocS-ΔAH was labeled with the red dye NT-647. Briefly, sample containing 50 nM of labeled RocS-ΔAH-6His and increasing concentrations of 6His-CpsC/D (from 275 pM to 9 µM) or ParB-6His (from 427 pM to 14 µM) were loaded on K023 Monolith NT.115 hydrophobic capillaries and thermophoresis was measured for 30 s at 25°C. Each measurement was made in triplicates. Experiments were carried out at 25°C in 10mM HEPES pH 7.5, 150mM NaCl and 0.05% Tween-20. Analysis was performed with the Monolith software. Affinity KD was quantified by analyzing the change in normalized fluorescence (Fnorm = fluorescence after thermophoresis/initial fluorescence) as a function of the concentration of the titrated 6His-CpsC/D or ParB-6His proteins.

### *oriC-ter* Ratio determination by real-Time qPCR

DNA genomic was extracted using the DNA maxima Kit (Qiagen). Real-time qPCR was performed as described previously (16). Briefly, each 20 µl sample consisted of 8.8 ng of DNA, 0.6 pmol of each primer (Table S2), and 10 µl of the 2x SYBR Green Supermix (Bio-Rad). Amplification was performed on an iQ5 Real-Time PCR Detection System (Bio-Rad). To find amplification efficiencies, Monte Carlo simulations were performed in R. Average C_t_-values and their corresponding standard deviations were used to simulate 10,000 new sets of C_t_-values that were used to compute the amplification efficiencies for each set. From that population of possible efficiencies, averages and standard deviations were derived. Analysis of the real-time qPCR experiments for *oriC-ter* ratio determination was performed using the 2^-ΔΔCT^ method (35), with the important difference that the earlier found amplification efficiencies were used to determine the fold-change per cycle, instead of assuming it to equal 2. As a reference, cells with an assumed *oriC*-*ter* ratio of 1 were used. For that, a thermo-sensitive DnaA-mutant (M398T) was grown at 30°C until an OD_600_ of 0.05. Then, cells were transferred to non-permissive temperature (40°C) and incubated for 1 hour, followed by harvesting and isolation of chromosomal DNA. Uncertainties in *oriC-ter* ratios were also determined by Monte Carlo simulations.

### Bioinformatic analyses

For the phylogenetic analysis, homologues of RocS were retrieved using iterative BLASTP from BLAST package 2.2.6 against a local database containing 4466 prokaryotic complete proteomes retrieved from NCBI ftp (ftp://ftp.ncbi.nlm.nih.gov/). The Spr0895 amino acid sequence (NP_358489.1) was used as first seed. Protein sequences detected as homologues were aligned with MAFFT v7.123b (36) and used to build an HMM profile with HMMER v3.1b1 (37). The profile was then used to query the local database with HMMSEARCH from the HMMER package. Plasmidic sequences have been removed from the analysis. Phylogeny of *Lactobacillales* has been inferred from a supermatrix of ribosomal proteins. One strain per family was selected to represent each family in *Lactobacillales* and a sequence of one species of *Listeriaceae* was added to root the tree. The sequences were aligned using MAFFT (L-INS-I option) and trimmed with BMGE-1.1 (option BLOSUM30) (38). The evolution model was chosen using BIC criteria and the phylogeny was inferred using PhyML (39) (LG+I+F+G4, 8 sequences, 6219 positions).

Secondary structure predictions of RocS were obtained using PSIPRED (40). The helical representation of RocS and MinD of *Escherichia coli* was made using http://www.tcdb.org/progs/?tool=pepwheel.

### Electrophoretic mobility shift assay (EMSA)

EMSA were carried out by incubating different concentrations of purified protein RocS-*Δ*AH-6His (0; 5; 10; 15 µM) with 50 ng of DNA in the following buffer (500mM Tris-HCl pH 8.8, 50mM MgSO_4_). DNA fragments of different length and percentage of GC content were PCR amplified (pUC18, *gfp* or genomic DNA of *Pseudomonas aeruginosa PA7*) using primers listed in Table S2. Reactions were incubated for 15 min at 37 °C before being loaded on 1% agarose gels. Gels were stained with ethidium bromide and revealed with UV light.

## Author Contributions

C.G. directed the study. C.M. conducted the experiments of cell biology, genetics with J.N., protein purification and western blot analysis. C.M. and A.D. performed image analyses. C.M. and N.D. implemented the Ori localization system. J.P.L. performed microscale thermophoresis experiments and contributed to protein purification. C.M. and J.S. performed *oriC*/*ter* ratio experiments. M.F.N.G. performed yeast two-hybrid experiments. P.S.G. performed phylogeny analyses. All authors designed and analyzed the data. C.G. and J.W.V. wrote the manuscript and all authors edited the manuscript. The authors declare no conflict of interest.

## Acknowledgments

Work on the Grangeasse lab is supported by grants from the CNRS, the University of Lyon, the Agence National de la Recherche (ANR-10-BLAN-1303-01 and ANR-15-CE32-0001-01), the Region Auvergne-Rhône-Alpes (financial support for C.M. and P.S.G.), the Fondation pour la Recherche Médicale (financial support for N.D. (ING20150532637) and C.M. (FDT20170437272)) and the Bettencourt-Schueller Foundation. Work in the Veening lab is supported by the Swiss National Science Foundation (project grant 31003A_172861), a VIDI fellowship (864.11.012) of the Netherlands Organization for Scientific Research (NWO-ALW), a JPIAMR grant (50-52900-98-202) from the Netherlands Organization for Health Research and Development (ZonMW) and ERC starting grant 337399-PneumoCell. We thank Stéphanie Ravaud for help in RocS structural predictions and Keith Weaver (University of South Dakota) for providing us with the pAD1 plasmid. We acknowledge the contribution of the Protein Science facility of the "SFR Biosciences Gerland-Lyon Sud (UMS344/US8)”.

**This article contains supporting information online.**

